# UNC-2 CaV2 channel localization at presynaptic active zones depends on UNC-10/RIM and SYD-2/Liprin-α in *Caenorhabditis elegans*

**DOI:** 10.1101/2021.01.27.428454

**Authors:** Kelly H. Oh, Mia Krout, Janet E. Richmond, Hongkyun Kim

## Abstract

Presynaptic active zone proteins couple calcium influx with synaptic vesicle exocytosis. However, the control of presynaptic calcium channel clustering by active zone proteins is not completely understood. In a *C. elegans* forward genetic screen, we find that UNC-10/RIM (Rab3-interacting molecule) and SYD-2/Liprin-*α* regulate presynaptic clustering of UNC-2, the CaV2 channel ortholog. We further quantitatively analyzed live animals using endogenously GFP-tagged UNC-2 and active zone components. Consistent with the interaction between RIM and CaV2 in mammals, the intensity and number of UNC-2 channel clusters at presynaptic terminals were greatly reduced in *unc-10* mutant animals. To understand how SYD-2 regulates presynaptic UNC-2 channel clustering, we analyzed presynaptic localization of endogenous SYD-2, UNC-10, RIMB-1/RIM-BP (RIM binding protein), and ELKS-1. Our analysis revealed that while SYD-2 is the most critical for active zone assembly, loss of SYD-2 function does not completely abolish presynaptic localization of UNC-10, RIMB-1, and ELKS-1, suggesting an existence of SYD-2-independent active zone assembly. UNC-2 localization analysis in double and triple mutants of active zone components show that SYD-2 promotes UNC-2 clustering by partially controlling UNC-10 localization, and ELKS-1 and RIMB-1 also contribute to UNC-2 channel clustering. In addition, we find that core active zone proteins are unequal in their abundance. While the abundance of UNC-10 at the active zone is comparable to UNC-2, SYD-2 and ELKS-1 are twice more and RIMB-1 four times more abundant than UNC-2. Together our data show that UNC-10, SYD-2, RIMB-1, and ELKS-1 control presynaptic UNC-2 channel clustering in redundant yet distinct manners.

**Significance Statement:** Precise control of neurotransmission is dependent on the tight coupling of the calcium influx through voltage-gated calcium channels (VGCCs) to the exocytosis machinery at the presynaptic active zones. However, how these VGCCs are tethered to the active zone is incompletely understood. To understand the mechanism of presynaptic VGCC localization, we performed a *C. elegans* forward genetic screen and quantitatively analyzed endogenous active zones and presynaptic VGCCs. In addition to RIM (Rab3-interacting molecule), our study finds that SYD-2/Liprin-*α* is critical for presynaptic localization of VGCCs. Yet, the loss of SYD-2, the master active zone scaffolding protein, does not completely abolish the presynaptic localization of the VGCC, showing that the active zone is a resilient structure assembled by redundant mechanisms.

## Introduction

The active zone of presynaptic terminals is a specialized compartment in which electrical signals are converted to chemical signals. In response to depolarization, voltage-gated calcium channels (VGCCs) deliver extracellular calcium ions, which in turn mediate the fusion of neurotransmitter-filled synaptic vesicles with the presynaptic membrane, resulting in neurotransmitter release and subsequent postsynaptic receptor activation (Sudhof, 2012; Dolphin and Lee, 2020). This conversion process requires close coupling of calcium influx with primed synaptic vesicles within the active zone. Thus, the precise positioning between docked synaptic vesicles and VGCCs is a critical determinant for the kinetics and efficiency of neurotransmitter release.

The organization of the active zone is structurally and/or functionally supported by a dense network of conserved scaffolding proteins of the active zone (CAZ) (Jin and Garner, 2008; Ackermann et al., 2015; Emperador-Melero and Kaeser, 2020). The core components of this scaffolding structure include Rab3-interaction protein (RIM), RIM-binding protein (RIM-BP), Liprin-*α*, and glutamine, leucine, lysine, and serine-rich protein (ELKS)/CAZ-associated structural protein (CAST). One intrinsic feature of active zone proteins is their multivalent interactions with other CAZ proteins. Distinct domains of single active zone proteins simultaneously mediate the interactions with other active zone proteins. Due to these redundant multivalent interactions, it is believed that the absence of a single active zone protein, with a few exceptions, does not alter the presynaptic localization of other active zone proteins or the structure of active zones (Acuna et al., 2016; Wang et al., 2016; Kushibiki et al., 2019).

The localization of the P/Q-type VGCC/CaV2.1, a major VGCC found at presynaptic active zones, is regulated by CAZ proteins. Previous studies have shown that the cytoplasmic region of the pore-forming CaV2.1 *α* _l_ subunit provides binding sites for RIM1/2, RIM-BP, and ELKS (Hibino et al., 2002; Fouquet et al., 2009; Kaeser et al., 2011; Kiyonaka et al., 2012). Complete ablation of all RIM proteins in mice reduces the level of CaV2.1 at active zones by approximately 50% (Kaeser et al., 2011). In contrast, removal of RIM-BP2 in mice does not reduce the overall levels of CaV2.1 and Ca^2+^ influx, but the relative positioning of CaV2.1 within active zones is altered (Acuna et al., 2015; Grauel et al., 2016). ELKS proteins have also been proposed to influence either CaV2 activation or levels at active zones but in a synapse-type specific manner (Liu et al., 2014; Held et al., 2016; Dong et al., 2018).

The role of individual active zone proteins in the presynaptic localization of synaptic vesicles, CaV2 channels, and other CAZ proteins has been assessed in loss-of-function mutants or genetically ablated invertebrate and mouse models. Because this strategy generally employs transgenic presynaptic markers or relies on immunostainings of synaptic proteins, it is often challenging to make a quantitative comparison. In this study, we took advantage of CRISPR/Cas9 genome editing to label endogenous key active zone proteins and CaV2/UNC-2 with GFP, and then assessed each of their *in vivo* synaptic localizations in *C. elegans* neurons. We found that the highly conserved RIM ortholog, UNC-10, plays a pivotal role in localizing the CaV2 channel ortholog, UNC-2, at presynaptic terminals. While the absence of ELKS-1 or RIMB-1 alone does not cause a significant alteration in presynaptic UNC-2 localization, the absence of both together with UNC-10 further reduces UNC-2 localization. Additionally, we found that SYD-2/Liprin-*α* is critical for UNC-2 presynaptic localization contributing via two distinct but overlapping mechanisms. SYD-2 maintains UNC-10 clustering at presynaptic terminals to indirectly regulate presynaptic UNC-2 clustering. SYD-2 also acts on other active zone proteins, including ELKS-1 and RIMB-1, to control presynaptic UNC-2 clustering.

## Results

### UNC-10/RIM and SYD-2/Liprin-*α* are required for presynaptic localization of UNC-2/CaV2

To understand how endogenous CaV2 channels are localized to presynaptic terminals, we inserted GFP into the amino-terminal region of UNC-2 in the endogenous locus using CRISPR/Cas9 genome editing (**Figure 1A**). The resulting animal, *cim104*[GFP::*unc-2*], was healthy and exhibited normal movements indistinguishable from wild-type animals. To analyze the functional integrity of endogenously tagged GFP::UNC-2, we performed a pharmacological assay using the acetylcholinesterase inhibitor aldicarb to detect alterations in *C. elegans* cholinergic synaptic transmission (Mahoney et al., 2006; Oh and Kim, 2017). Decreased sensitivity to aldicarb-mediated paralysis is indicative of reduced synaptic transmission. GFP-tagged UNC-2 *cim104* animals showed aldicarb sensitivity comparable to wild-type N2 animals (**Figure 1 supplement 1**). We also determined whether *cim104* animals exhibit normal evoked responses at neuromuscular junctions, using patch clamp electrophysiology and found that evoked responses were unperturbed compared to wild-type animals (**Figure 1 supplement 2**). To further ensure that GFP::UNC-2 localizes to presynaptic active zones, we examined UNC-2 co-localization with the presynaptic active zone protein RIMB-1, the ortholog of mammalian RIM binding protein 1 and 2. The coding sequence of the red fluorescent mScarlet protein was inserted into the endogenous *rimb-1* locus at its initiation codon. UNC-2 and RIMB-1 imaged in the dorsal nerve cord co-localized in all synaptic puncta (**Figure 1B**), demonstrating that the genomic tagging of UNC-2 and RIMB-1 does not interfere with their previously reported synaptic colocalization (Kurshan et al., 2018).

**Figure 1.**
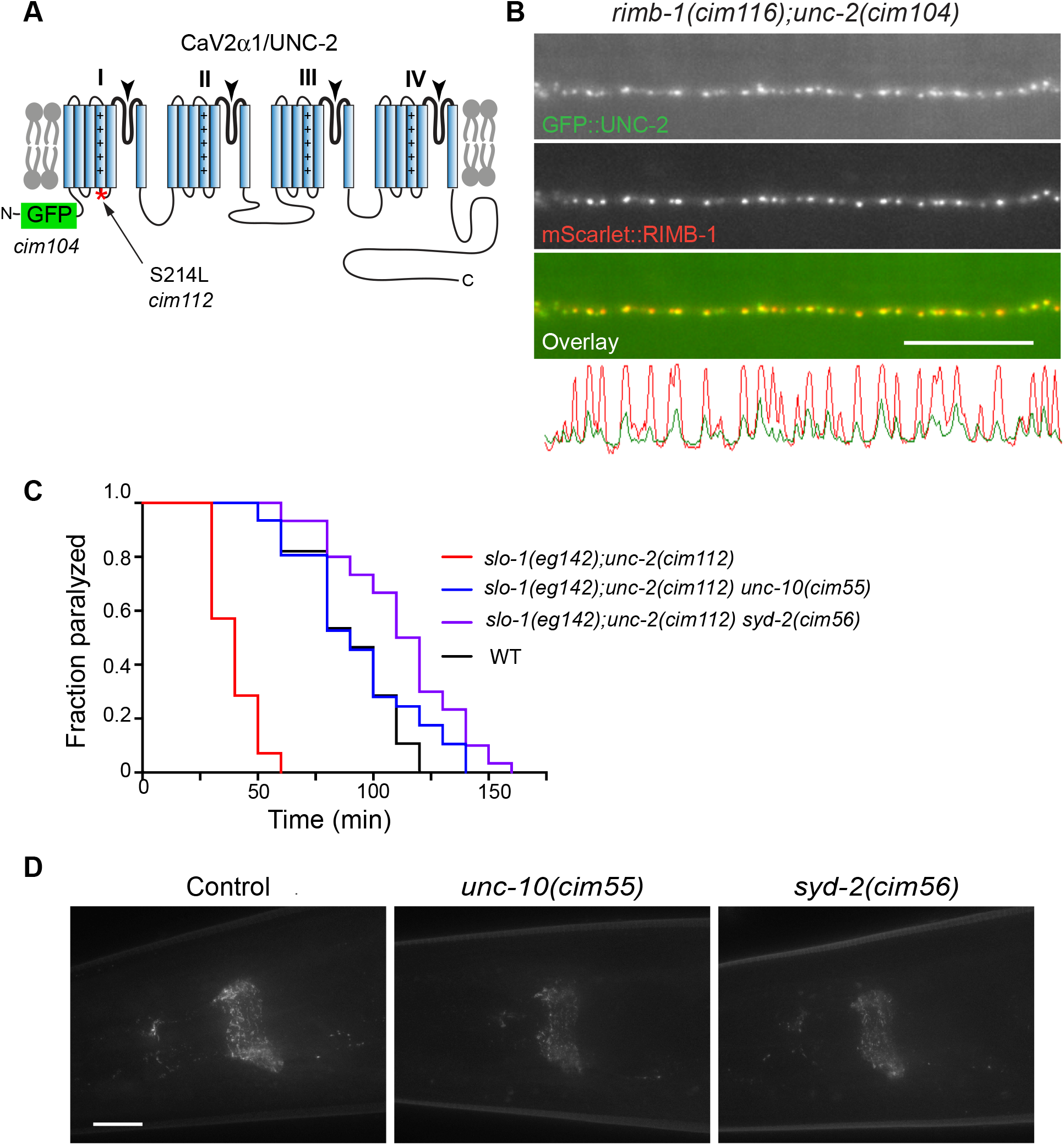
A genetic screen identifies *unc-10/*RIM and *syd-2/*Liprin-*α* as major genes required for UNC-2 localization at presynaptic terminals. (**A**) The predicted structure of UNC-2. The relative positions of CRISPR/Cas9-mediated GFP insertion and amino acid substitution (S218L) are denoted. (**B**) GFP::UNC-2 and mScarlet::RIMB-1 are co-localized at the dorsal nerve cord. Scale bar:10 μm (**C**) Two presynaptic mutants *unc-10(cim55)* and *syd-2(cim56)* suppress aldicarb hypersensitivity of *slo-1(eg142);unc-2(cim112[GFP::unc-2S218])* animals. p < 0.0001, Log-rank survival test. (**D**) UNC-2 images in the nerve rings of *slo-1(eg142);unc-2(cim112), slo-1(eg142);unc-2(cim112) unc-10(cim55)*, and *slo-1(eg142);unc-2(cim112) syd-2(cim56)* mutant animals. The presented images are maximum intensity projection images of a 40 Z-stack captured at 0.2 μm intervals. Scale bar:10 μm

To understand the molecular mechanism underlying CaV2 channel localization to the presynaptic active zone, we designed a genetic screen. We first generated the *cim112* animal, in which an S218L point mutation was introduced into the coding sequence of UNC-2 of the *cim104* animal (**Figure 1A**). This S218L mutation and the corresponding mutation in human CaV2 have been shown to enhance calcium channel function therefore providing a sensitized background for the screen (van den Maagdenberg et al., 2007; Huang et al., 2019). Because the BK channel ortholog SLO-1 can modulate the function of UNC-2 channels making it difficult to interpret the nature of genetic interactions with genes obtained from the screen (Oh et al., 2015; Oh et al., 2017), we introduced a *slo-1* null mutation, *eg142*, into the *cim112* animal. As expected, the combined effects of *slo-1(eg142);unc-2(cim112[GFP::unc-2(S218L)])* mutations produced animals that were aldicarb-hypersensitive (**Figure 1C, Figure 1-supplement 1**). In this background we screened for mutants that showed resistance to the paralyzing effects of aldicarb in F2 progeny of chemically mutagenized *slo-1(eg142);unc-2(cim112)* double mutant animals (**Figure 1 supplement 3**).

This screen proved highly selective in identifying mutants that disturb *unc-2* function. In addition to *unc-2* loss- or reduction-of-function mutants, the screen yielded multiple alleles of *unc-36* and *calf-1*, which encode a CaV2 *α* _2_*δ* subunit ortholog and an ER membrane protein required for UNC-2 trafficking, respectively (Saheki and Bargmann, 2009). Importantly, two *unc-10* and one *syd-2* mutants were also identified in the screen, although other mutants that have a defect in CAZ have not been identified thus far. Compared to wild type animals, both *unc-10(cim55)* and *syd-2(cim56)* animals show significantly decreased GFP-tagged UNC-2 levels in the nerve ring, a collection of synapses derived from over half of the entire neurons (**Figure 1D**). Based on these results, we more precisely determined how UNC-10 and SYD-2 are involved in UNC-2 channel cluster formation at presynaptic terminals.

For quantification of UNC-2 channel clusters at presynaptic terminals, we used the *unc-10(md1117)* and *syd-2(ok217)* null mutant strains. In *C. elegans* as well as *Drosophila*, and mammals, UNC-10 and/or SYD-2 homologs are known to interact with their corresponding RIMB-1 and ELKS-1 homologs. Furthermore, mammalian RIM-BP2 and ELKS proteins have been reported to play a role in CaV2 localization in certain contexts (Liu et al., 2014; Acuna et al., 2015; Grauel et al., 2016; Held et al., 2016; Dong et al., 2018). Therefore, we included *rimb-1(ce828)* and *elks-1(ok2762)* mutants in our analysis of UNC-2 channel clustering. For quantitative comparison, we determined the fluorescence intensity of clustered GFP-tagged UNC-2 puncta in a select posterior area of the dorsal nerve cord, where cholinergic and GABAergic neurons make discrete *en passant* synapses with the dorsal body wall muscles (**Figure 2-supplement 1**). We found that the number and intensity of GFP::UNC-2 clusters were greatly reduced in *unc-10(md1117)* compared to wild-type animals (**Figure 2**). This finding is consistent with a previous study from conditional knock-out mice of all RIM isoforms (Kaeser et al., 2011). *syd-2(ok217)* mutants showed an even more pronounced reduction in UNC-2 cluster number and intensity than *unc-10(md1117)* mutants. In contrast, *rimb-1(ce828)* and *elks-1(ok2762)* mutants were comparable to wild-type animals in number and intensity of UNC-2 clusters

**Figure 2.**
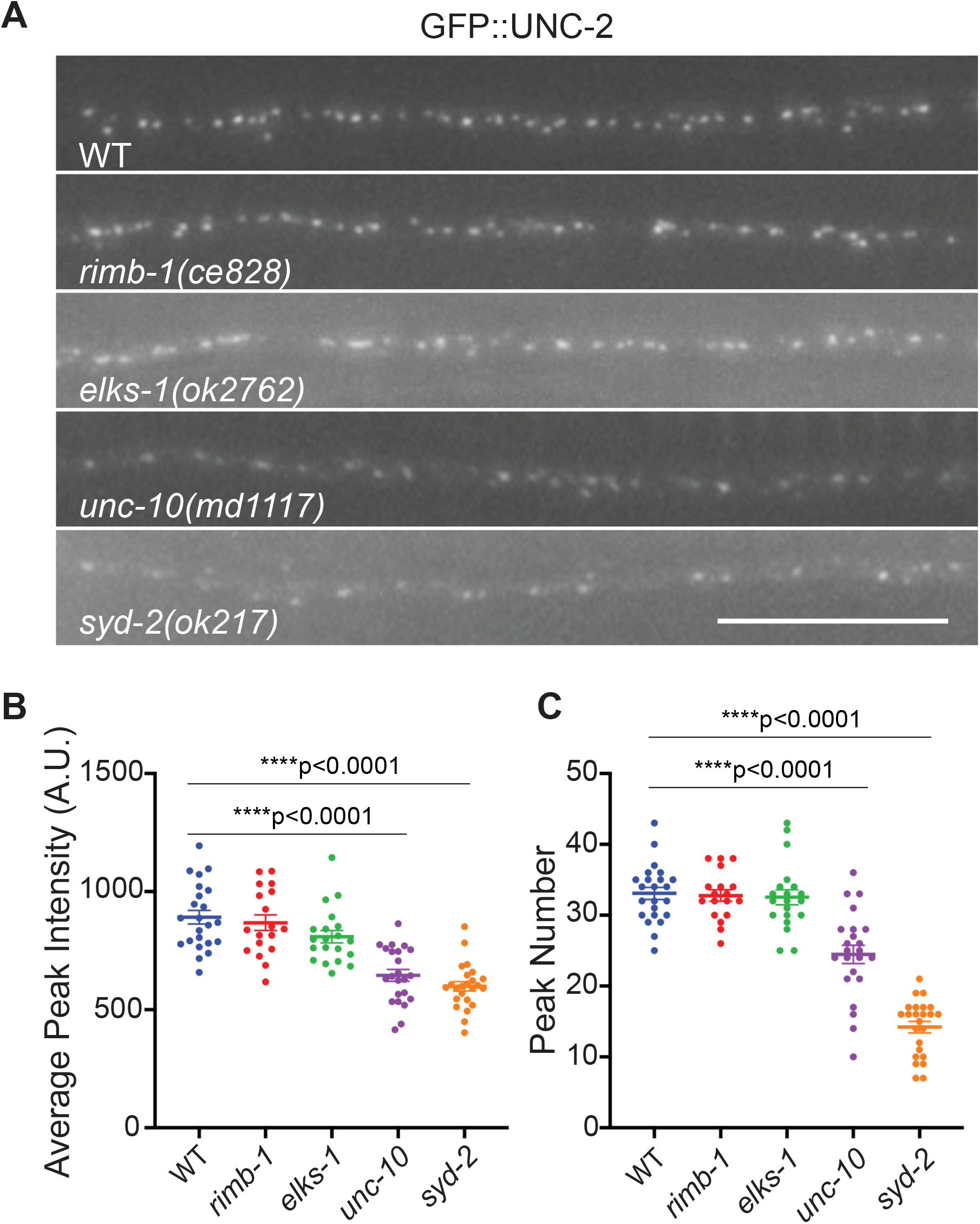
UNC-2 channel clustering at presynaptic terminals is regulated by UNC-10 and SYD-2. (**A**) Endogenous UNC-2 clustering in motor neurons of the posterior dorsal nerve cord is reduced in *unc-10* and *syd-2* mutants, but not in *elks-1* and *rimb-1* mutants. Scale bar, 10 μm. (**B**) Average UNC-2 peak intensity in wild-type (WT), *rimb-1, elks-1, unc-10*, and *syd-2* mutant animals. (**C**) The number of UNC-2 clusters in a 30 μm interval of wild-type (WT), *rimb-1, elks-1, unc-10*, and *syd-2* mutant dorsal nerve cords. Error bars represent SEM. One-way ANOVA with Tukey’s post-hoc analysis.

### SYD-2 acts as a master scaffolding protein for presynaptic localization of UNC-10, RIMB-1, and ELKS-1

The results above demonstrate that SYD-2 and UNC-10 impact UNC-2 puncta to differing degrees. SYD-2 has been reported to act as a master synapse assembly protein in *C. elegans* and prior studies have demonstrated interactions with RIM proteins as well as other CAZ components including RIM-BP and ELKS (Ohtsuka et al., 2002; Wang et al., 2002; Ko et al., 2003; Dai et al., 2006; Kaeser et al., 2011; Petzoldt et al., 2020). Therefore, we considered the possibility that the effects of the *syd-2* mutation on UNC-2 could be due to synaptic alterations of each of these binding partners. To test this possibility, we generated endogenously GFP-tagged strains of UNC-10, RIMB-1, and ELKS-1, as well as SYD-2 using CRISPR/Cas9 genome editing to examine the interdependence of these active zone proteins for synaptic localization. Unlike synaptic vesicle markers that show faint signals along the axon while enriched within presynaptic terminals (Oh et al., 2015), these GFP-tagged active zone proteins showed more clear, discrete puncta.

We first compared the peak intensities and numbers of UNC-10 presynaptic clusters in wild-type, *rimb-1(ce828), elks-1(ok2762)*, and *syd-2(ok217)* animals (**Figure 3**). The peak intensity and number of presynaptic UNC-10 clusters were reduced in *syd-2(ok217)* mutant animals, but not in *rimb-1(ce828)* or *elks-1(ok2762)* mutant animals. Having observed a reduction in UNC-10 clustering in *syd-2(ok217)* mutant animals, we then performed the reciprocal experiments to determine whether SYD-2 presynaptic clusters are altered in *unc-10(md1117)* mutants as well as in *rimb-1(ce828)* or *elks-1(ok2762)* mutants. The peak intensities of SYD-2 clusters were unaffected in all three mutants, confirming that SYD-2 is the master regulator of the presynaptic CAZ organization (Zhen and Jin, 1999; Dai et al., 2006; Patel et al., 2006). These results together, indicate that the reduction of UNC-2 clusters in *syd-2(ok217)* mutants can be partially explained by the role of SYD-2 in UNC-10 presynaptic localization (**Figure 3**). Since the *syd-2* mutation did not completely abolish presynaptic UNC-10 localization, UNC-2 clustering could be supported by the remaining UNC-10 proteins at presynaptic terminals. In this case, we predicted that UNC-2 clustering would be more severely affected in *unc-10(md1117)* mutant animals than *syd-2(ok217)* mutant animals. In contrast to this prediction, we observed that *syd-2(ok217)* mutant animals exhibit a more severe defect in UNC-2 clustering than *unc-10(md1117)* mutant animals (**Figure 2**). Thus, it is likely that SYD-2 provides an additional mechanism for UNC-2 clustering, independent of UNC-10 localization.

**Figure 3.**
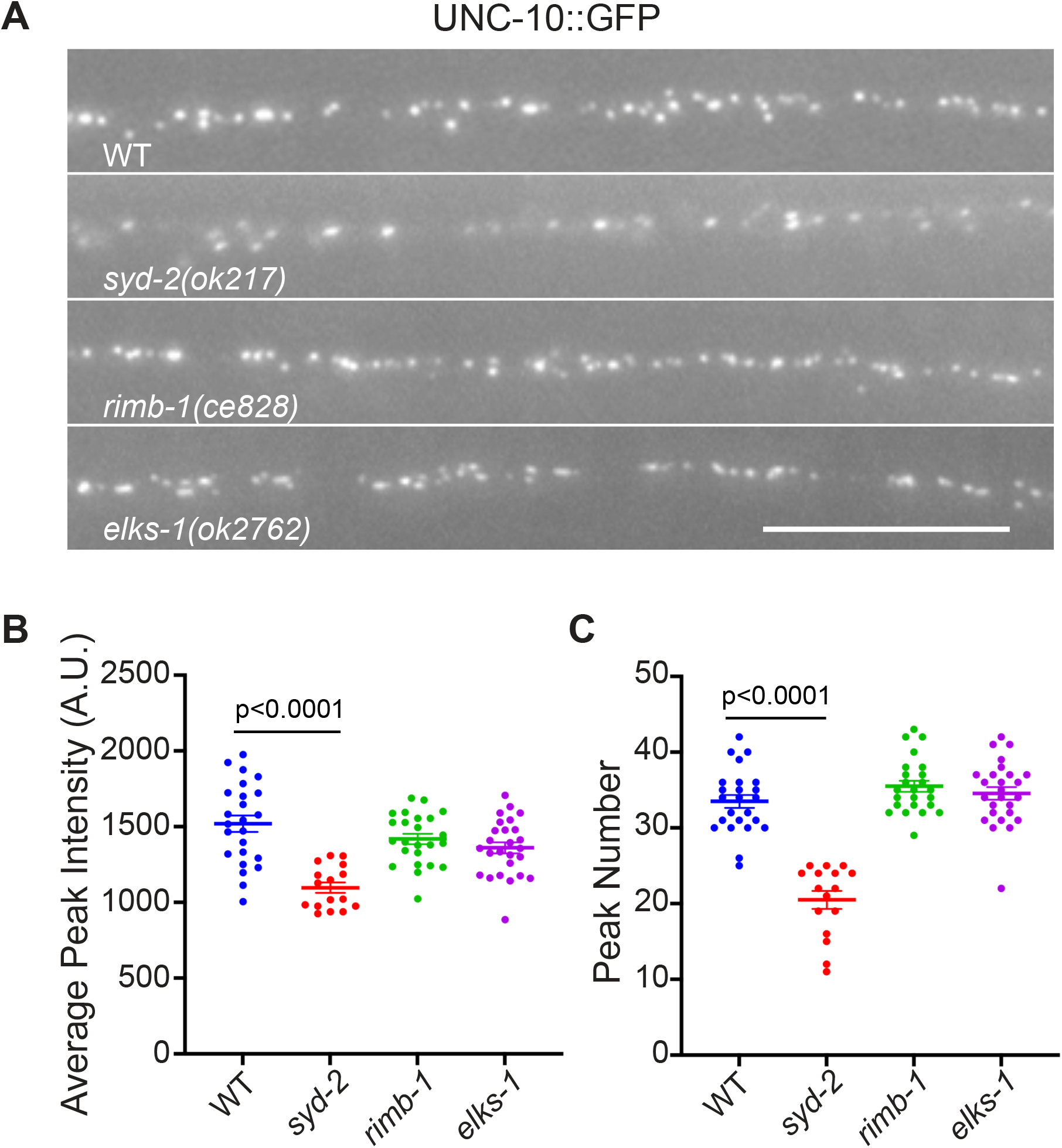
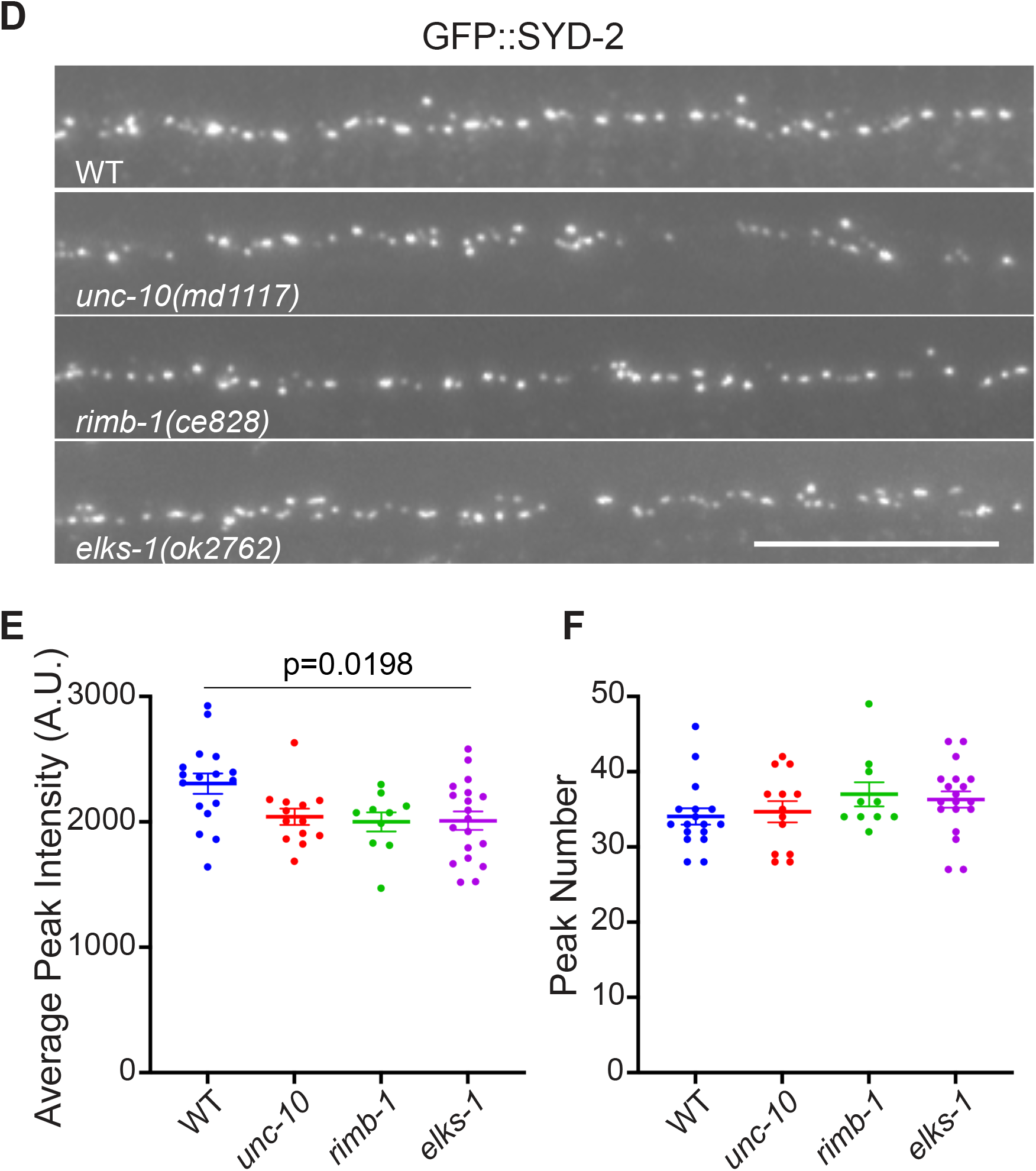
SYD-2 is a scaffolding protein upstream of UNC-10, RIMB-1, and ELKS-1. (**A**) UNC-10 clusters at presynaptic terminals are reduced, but not eliminated, in *syd-2* mutant animals. Endogenous UNC-10 clustering in motor neurons of the posterior dorsal nerve cord is reduced in *syd-2* mutants, but not in *elks-1* and *rimb-1* mutants. Scale bar, 10 μm. (**B**) Average UNC-10 peak intensity in wild-type (WT), *rimb-1, elks-1*, and *syd-2* mutant animals. (**C**) The number of UNC-10 clusters in the dorsal nerve cords (per a 30 μm interval) of wild-type (WT), *rimb-1, elks-1*, and *syd-2* mutant animals. (**D**) Endogenous SYD-2 clustering in motor neurons of the posterior dorsal nerve cord is largely independent of *unc-10, elks-1, rimb-1* mutants. Scale bar, 10 μm. (**E**) Average SYD-2 peak intensity in wild-type (WT), *rimb-1, elks-1*, and *unc-10* mutant animals. (**F**) The number of SYD-2 clusters in the dorsal nerve cords (per a 30 μm interval) of wild-type (WT), *rimb-1, elks-1*, and *unc-10* mutant animals. Error bars represent SEM. One-way ANOVA with Tukey’s post-hoc analysis.

Given that SYD-2 associates with and recruits multiple CAZ proteins, we next explored the possibility that in addition to UNC-10, SYD-2 organizes UNC-2 channel clusters at presynaptic terminals through the recruitment of RIMB-1 and EKLS-1. First, we examined if RIMB-1 and ELKS-1 localization are dependent on SYD-2. While the intensities of RIMB-1 clusters are reduced in *unc-10(md1117), elks-1(ok2762)*, and *syd-2(ok217)* mutant animals, the extent of the reduction was most prominent in *syd-2(ok217)* mutant animals. In contrast to RIMB-1 puncta intensities, the number of RIMB-1 clusters was only affected in *syd-2(ok217)* mutants, and was accompanied by diffuse intrasynaptic RIMB-1 protein expression (**Figure 4**). Likewise the GFP intensities of ELKS-1 clusters were also reduced in *unc-10(md1117), rimb-1(ce828)*, and *syd-2(ok217)* mutant animals, with a reduction in synapse number only apparent in *syd-2(ok217)* mutant animals (**Figure 4**). These results indicate that SYD-2 is the main active zone protein responsible for recruiting or stabilizing UNC-10, RIMB-1, and ELKS-1 at presynaptic active zones, while these three proteins also contribute to active zone assembly or stabilization.

**Figure 4.**
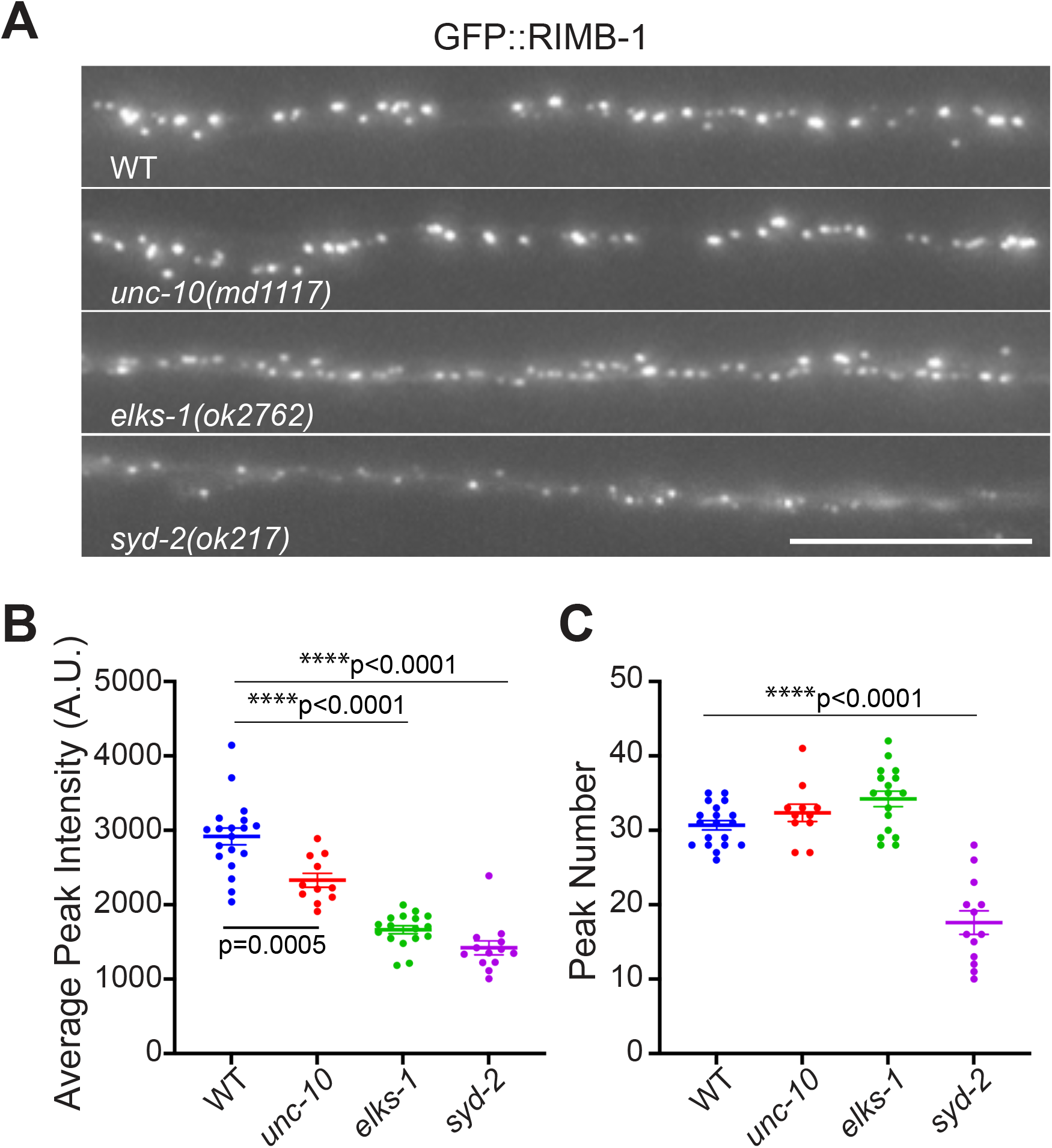
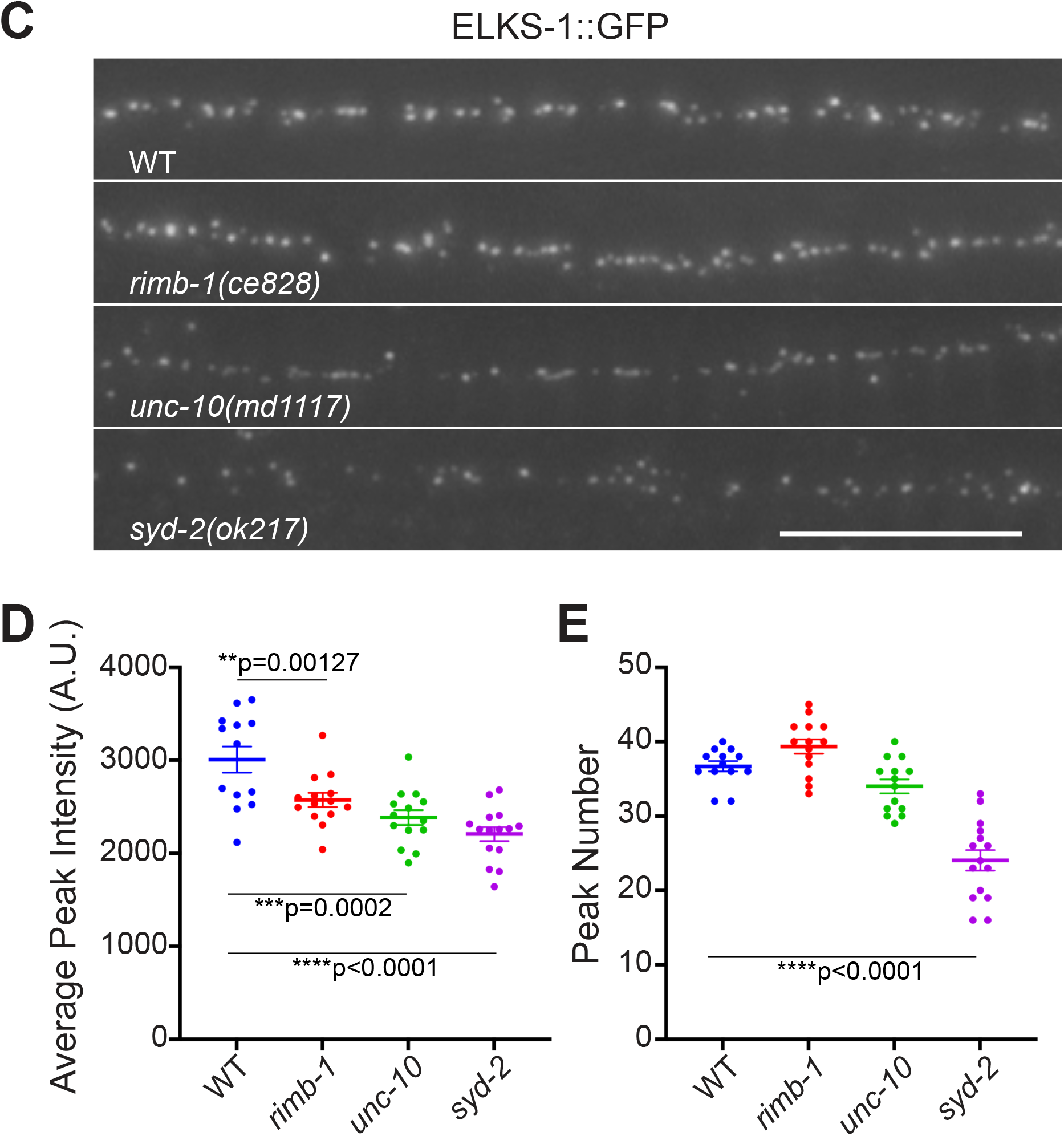
The localization of RIMB-1 and ELKS-1 at presynaptic terminals is partially dependent on SYD-2. (**A**) The number and intensity of RIMB-1 clusters are reduced, but not eliminated, in *syd-2* mutant animals, and UNC-10 and ELKS-1 are required for the normal level of cluster intensity. Scale bar, 10 μm. (**B**) Average RIMB-1 peak intensity in wild-type (WT), *rimb-1, elks-1*, and *syd-2* mutant animals. (**C**) The number of RIMB-1 clusters in the dorsal nerve cords (per 30 μm) of wild-type (WT), *unc-10, elks-1*, and *syd-2* mutant animals. (**D**) Endogenous ELKS-1 clustering in motor neurons of the posterior dorsal nerve cord largely depends on SYD-2. Scale bar, 10 μm. (**E**) Average ELKS-1 peak intensity in wild-type (WT), *rimb-1, elks-1*, and *unc-10* mutant animals. (**F**) The number of ELKS-1 clusters in the dorsal nerve cords (per 30 μm) of wild-type (WT), *rimb-1, elks-1*, and *unc-10* mutant animals. Error bars represent SEM. ****p<0.0001. One-way ANOVA with Tukey’s post-hoc analysis.

### Stoichiometry of UNC-2, UNC-10, RIMB-1, ELKS-1, and SYD-2

In imaging endogenously labelled CAZ proteins, we noticed that relative presynaptic GFP fluorescence intensities were markedly different for the components examined. To quantify these relative levels we imaged using a camera with a higher dynamic range (**Figure 5**). We found that the average and maximum intensities of UNC-2/CaV2and UNC-10/RIM puncta were comparable at presynaptic terminals. In contrast, the average and maximum intensities of presynaptic SYD-2 and ELKS-1 puncta were significantly higher than UNC-2 and UNC-10 puncta by a factor of 2. Whether this two-fold increase in SYD-2 and ELKS-1 label intensity relates to previously reported observations that SYD-2 forms homodimers at presynaptic terminals (Taru and Jin, 2011) and physically interact with ELKS-1 (Deken et al., 2005) remains to be tested. Unexpectedly, RIMB-1 is four times more enriched than UNC-2 or UNC-10 at synapses. The stoichiometry of presynaptic CaV2 and CAZ proteins may reflect differences in copy number due to homodimerization and multimeric assembly of some components.

**Figure 5.**
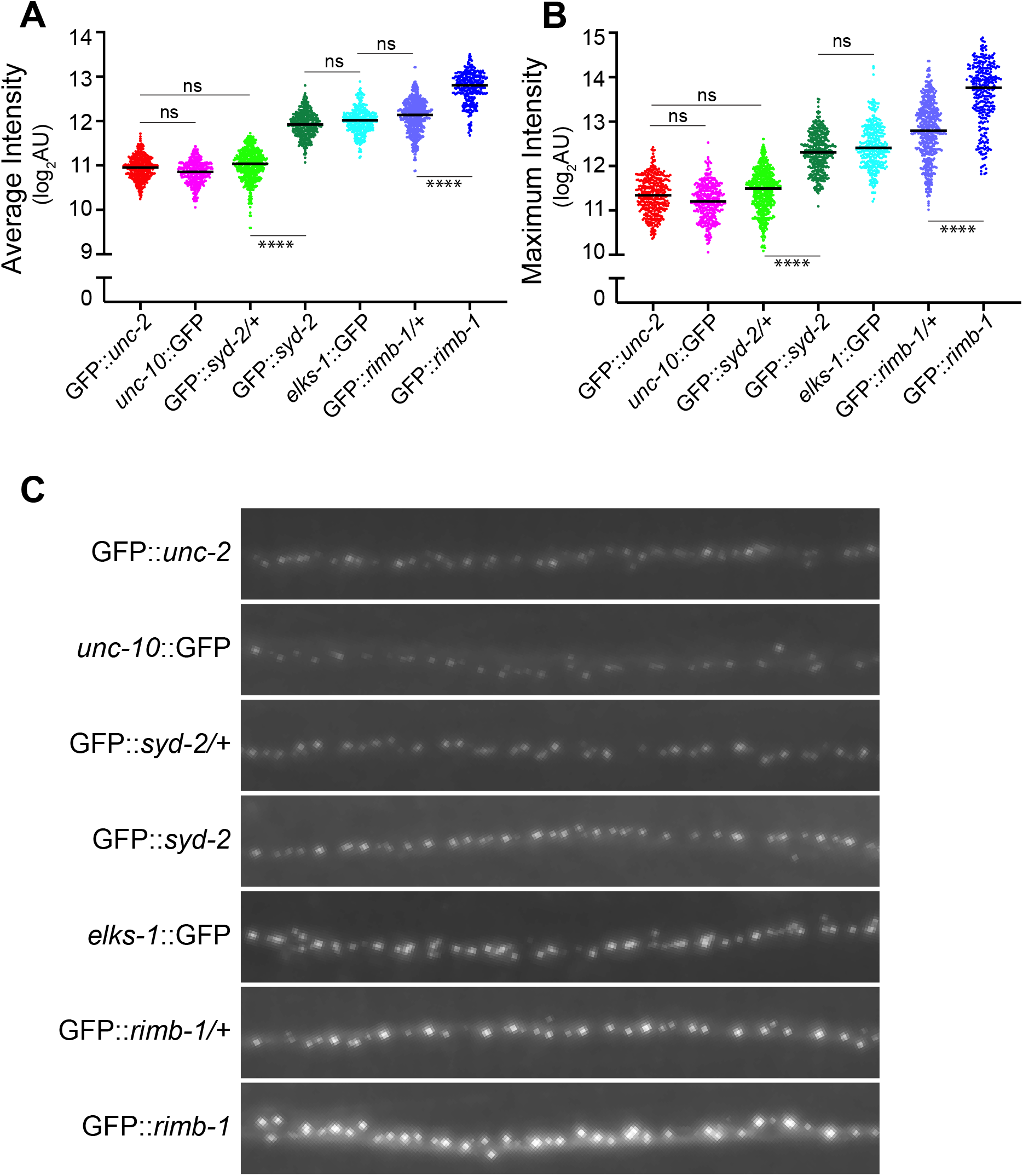
Copy number analysis of UNC-2/CaV2 and cytomatrix proteins at the presynaptic terminals indicates at least three distinct groups. The relative levels of endogenously GFP-tagged UNC-2, UNC-10, SYD-2, ELKS-1, and RIMB-1 at the presynaptic terminals are determined in the dorsal cord of animals. To evaluate the relative intensity of GFP puncta, heterozygous GFP::*syd-2*/+ and GFP::*rimb-1*/+ animals were also included in the analysis. (**A**) The average intensity of UNC-2, UNC-10, SYD-2/+, SYD-2, ELKS-1, RIMB-1/+, and RIMB-1 puncta. The data do not pass a normality test, and Kruskal-Wallis test with Dunn’s post-hoc was used for statistical analysis (p<0.0001). (**B**) The maximum intensity of UNC-2, UNC-10, SYD-2/+, SYD-2, ELKS-1, RIMB-1/+, and RIMB-1 puncta. The number of puncta ranges approximately from 300 to 600, as all of the focused puncta within the visual field were pooled from 10 independent animals. ****p<0.0001, Kruskal-Wallis test with Dunn’s post-hoc analysis. (**C**) The representative images of UNC-2, UNC-10, SYD-2/+, SYD-2, ELKS-1, RIMB-1/+, and RIMB-1 puncta. Images were scaled to the same dynamic range to visualize puncta.

### RIMB-1 and ELKS-1 contribute to synaptic localization of the UNC-2/CaV channel

RIM, RIM-BP, and ELKS have been previously shown to interact with CaV2 channels, while SYD-2/Liprin-*α* has not been implicated in CaV2 channel localization. It is possible that SYD-2 can directly act on UNC-10 and UNC-10 subsequently recruits RIMB-1 or ELKS-1 to form UNC-2 channel cluster formation at presynaptic terminals. Alternatively, SYD-2 may act on RIMB-1 or ELKS-1 independently of UNC-10 to regulate UNC-2 channel cluster formation. To test these possibilities, we first sought to understand the relationship between UNC-10 with RIMB-1 and ELKS-1 by comparing GFP::UNC-2 clusters in *unc-10(md1117)* single, double, and triple mutants with *rimb-1(ce828)* and *elks-1(ok2762)* (**Figure 6**). While the absence of ELKS-1 did not further alter UNC-2 clusters, the absence of RIMB-1 moderately reduced the peak intensity of UNC-2 clusters of *unc-10(md1117)* mutant animals without significantly changing the overall number of UNC-2 clusters. These results indicate that RIMB-1, but not ELKS-1, contributes to UNC-2 localization when UNC-10 is absent. Interestingly, *unc-10(md1117);rimb-1(ce828)*;*elks-1(ok2762)* triple mutant animals exhibit a further reduction in both number and peak intensity of UNC-2 clusters compared to the *unc-10(md1117);rimb-1(ce828)* double mutant animals. Thus, RIMB-1 and ELKS-1 also contribute to UNC-2 localization, although UNC-10 plays the predominant role.

**Figure 6.**
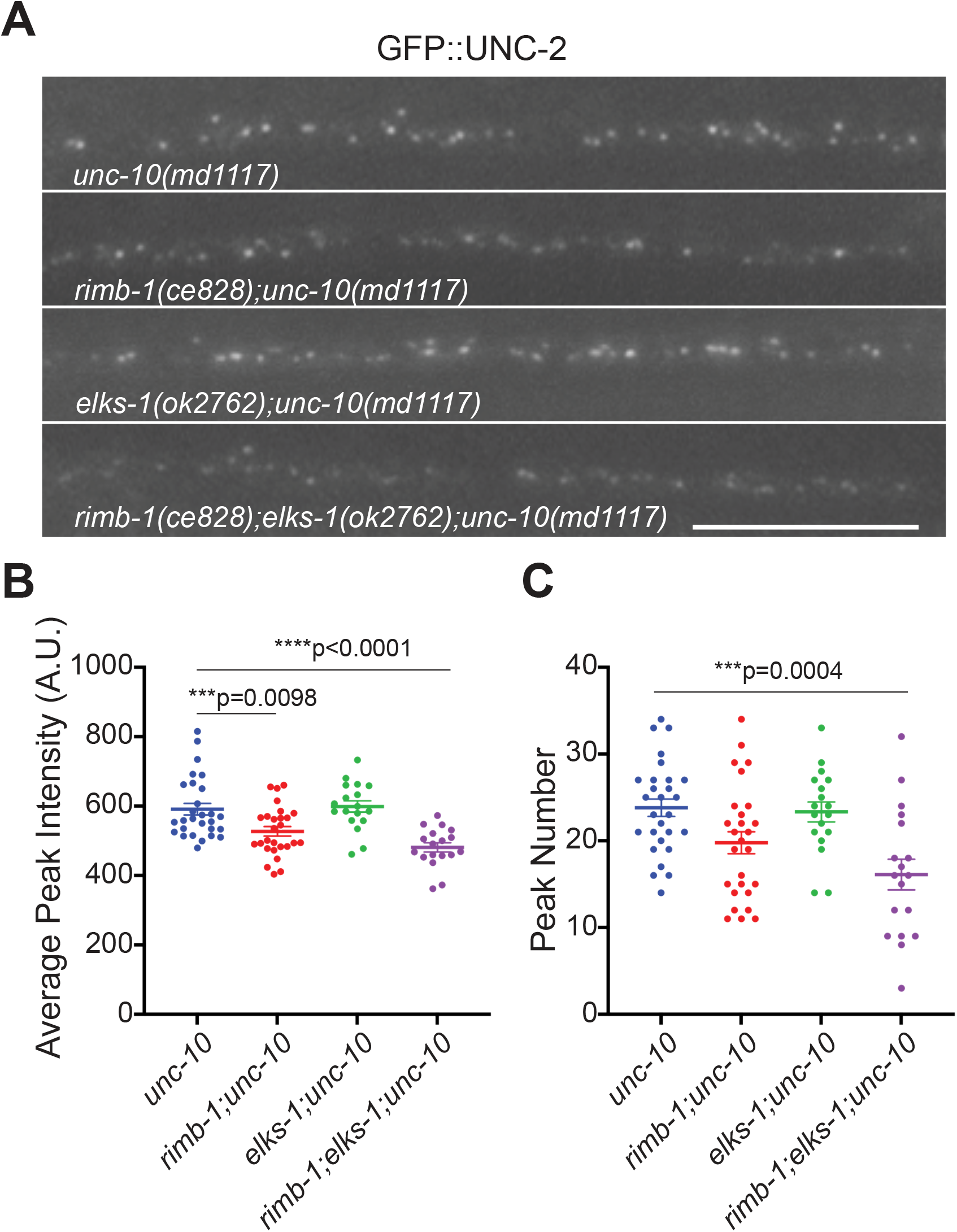
RIMB-1 and ELKS-1 contribute to UNC-2 cluster formation. (**A**) Endogenous UNC-2 clustering in motor neurons of the posterior dorsal nerve cord is further reduced in rimb-1;*unc-10* double and rimb-1;elks-1;*unc-10* triple mutants. Scale bar, 10 μm. (**B**) Average UNC-2 peak intensity in *unc-10, rimb-1;unc-10, elks-1;unc-10*, and *rimb-1;elks-1;unc-10* mutant animals. One-way ANOVA with Tukey’s post-hoc analysis. (**C**) The number of UNC-2 clusters in the dorsal nerve cords (per 30 μm) of *unc-10, rimb-1;unc-10, elks-1;unc-10*, and *rimb-1;elks-1;unc-10* mutant animals.

### UNC-2/CaV is localized at the synapse by both SYD-2-dependent and -independent mechanisms

Next, we investigated how SYD-2 regulates UNC-2 channel clustering at presynaptic terminals in relation to UNC-10, ELKS-1, and RIMB-1. Although the localization of UNC-10, RIMB-1, and ELKS-1 depends largely on SYD-2, all three proteins show residual localization in the absence of SYD-2. To determine whether the remaining GFP::UNC-2 in *syd-2* mutants is due to the residual UNC-10, RIMB-1, and ELKS-1, we compared UNC-2 clusters in *unc-10(md1117) syd-2(ok217), elks-1(ok2762);syd-2(ok217)*, and *rimb-1(ce828);syd-2(ok217)* double mutant animals (**Figure 7**). Compared to *syd-2(ok217)* single mutant animals, presynaptic UNC-2 was further reduced in *unc-10(md1117) syd-2(ok217)* and *elks-1(ok2762);syd-2(ok217)* mutant, but not in *rimb-1(ce828);syd-2(ok217)* mutant. This suggests that SYD-2-independent UNC-10 and ELKS-1 at active zones contributes to UNC-2 localization, while the contribution of RIMB-1 is largely dependent on SYD-2 (**Figure 7-supplement 1**). Next, we examined whether the three proteins, UNC-10, RIMB-1, and ELKS-1, together can account for presynaptic UNC-2 clustering. Surprisingly, UNC-2 clusters are still present in *rimb-1(ce828);elks-1(ok2762);unc-10(md1117)* triple mutants, although their number and intensity are slightly lower than in *syd-2(ok217)* mutants. This indicates that SYD-2 can localize UNC-2 either through a direct interaction or through a protein other than UNC-10, RIMB-1, or ELKS-1.

**Figure 7.**
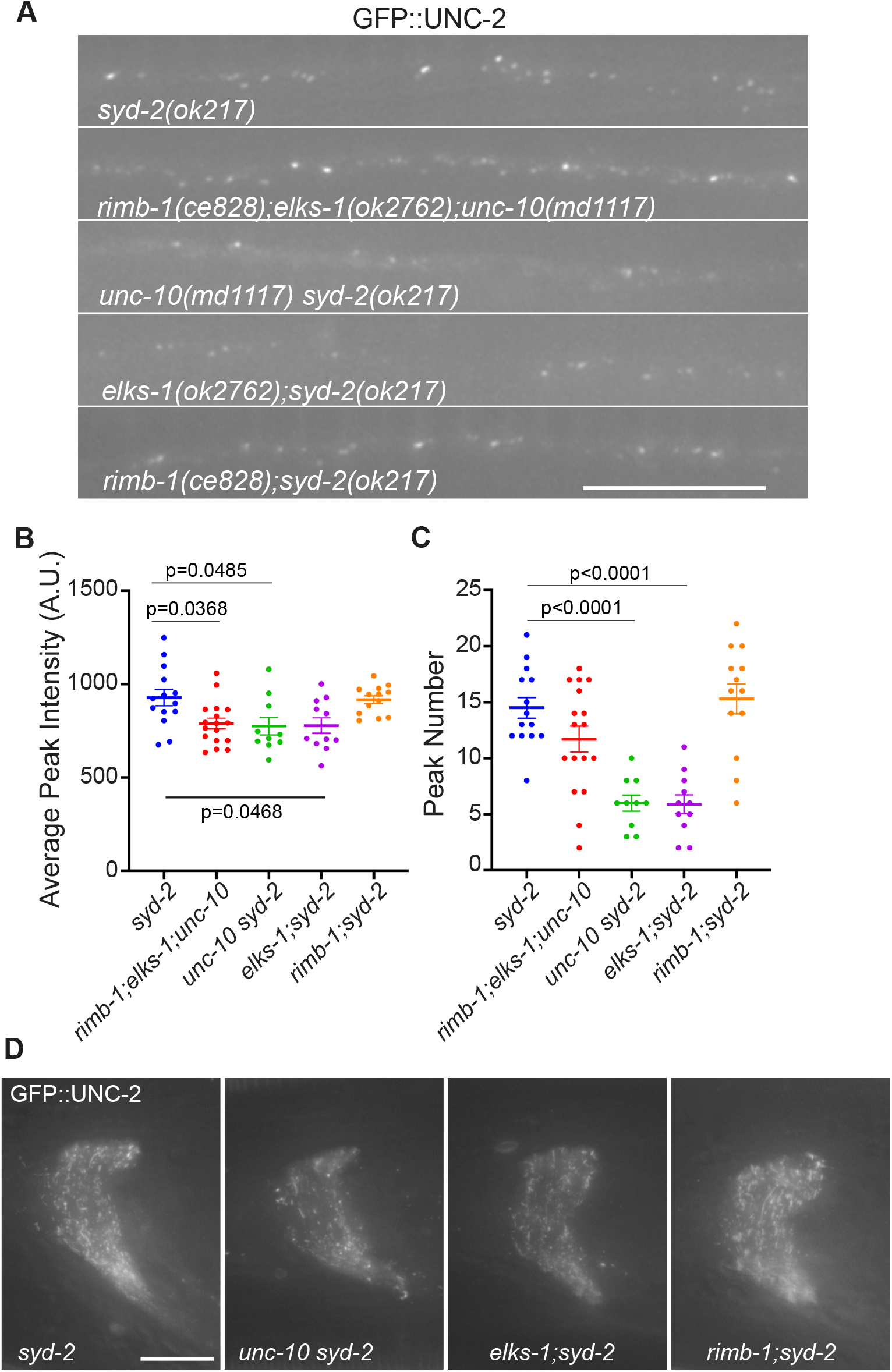
UNC-2 clustering is dependent on both SYD-2-dependent and -independent mechanisms. (**A**) Endogenous UNC-2 clustering in motor neurons of the posterior dorsal nerve cord is further reduced in *unc-10 syd-2* and *elsk-1;syd-2* double mutants. Scale bar, 10 μm. (**B**) Average UNC-2 peak intensity in *syd-2, rimb-1;elks-1;unc-10, unc-10 syd-2, elks-1;syd-2, rimb-1;syd-2* mutant animals. Error bars represent SEM. ****p<0.0001. One-way ANOVA with Tukey’s post-hoc analysis. (**C**) The number of UNC-2 clusters in the dorsal nerve cords (per 30 μm) of *syd-2, rimb-1;elks-1;unc-10, unc-10 syd-2, elks-1;syd-2, rimb-1;syd-2* mutant animals. Error bars represent SEM. ****p<0.0001. One-way ANOVA with Tukey’s post-hoc analysis. (**D**) UNC-2 images in the nerve rings of *syd-2, unc-10 syd-2, elks-1;syd-2*, and *rimb-1;syd-2* mutant animals. The images are maximum intensity projection images of a 45 Z-stack captured at 0.2 μm intervals. A reduction of UNC-2 in *unc-10 syd-2* and *elks-1;syd-2* mutant animals is prominent compared to *syd-2* and *rimb-1;syd-2* mutant animals. Scale bar, 10 μm.

### UNC-104/KIF1A is required for axonal transport of SYD-2, UNC-10, RIMB-1, ELKS-1, and UNC-2

In addition to synapse assembly/stability, SYD-2 has been reported to have a role in the transport of synaptic vesicles and active zone proteins (Wagner et al., 2009; Wu et al., 2013; Edwards et al., 2015). Thus, it could be argued that the role of SYD-2 in presynaptic UNC-2 clustering can be entirely explained by its upstream role in the axonal transport of CAZ proteins. UNC-104/KIF1A is the major kinesin motor protein necessary for the transport of synaptic vesicles (Hall and Hedgecock, 1991; Okada et al., 1995). We examined if UNC-104 is necessary for the synaptic localization of UNC-2, UNC-10, RIMB-1, ELKS-1, and SYD-2 in the dorsal nerve cord. Since our results indicate that SYD-2 plays a key role in the active zone localization of UNC-2, UNC-10, RIMB-1, and ELKS-1, we first examined SYD-2 localization in a hypomorphic allele of *unc-104(e1265)* mutant known to severely impair vesicle transport while avoiding the lethality associated with *unc-104* null mutants (**Figure 8**). SYD-2 expression was significantly reduced in the dorsal nerve cord, indicating that axonal transport of SYD-2 is dependent on UNC-104. Since UNC-2, UNC-10, RIMB-1, and ELKS-1 are partially dependent on SYD-2 for their active zone localization, their synaptic expression was also expected to be disrupted in *unc-104* mutant. Indeed, the dorsal nerve cord localization of all four proteins was disrupted in *unc-104* mutant (**Figure 8**). In addition, the number of UNC-2, UNC-10, and ELKS-1 puncta is consistently lower in *unc-104* mutants than *syd-2* mutants. Unlike UNC-2, UNC-10, and ELKS-1, RIMB-1 puncta are highly variable in size and number and mobile in *unc-104* mutants (**Video 1**). Together, the localization of UNC-2, UNC-10, RIMB-1, and ELKS-1 was more severely disrupted in *unc-104* mutants than in the *syd-2* mutant background. This is consistent with the observations that a fraction of UNC-10, RIMB-1, and ELKS-1 proteins can localize to presynaptic terminals independently of SYD-2, and this SYD-2-independent localization is dependent on UNC-104.

**Figure 8.**
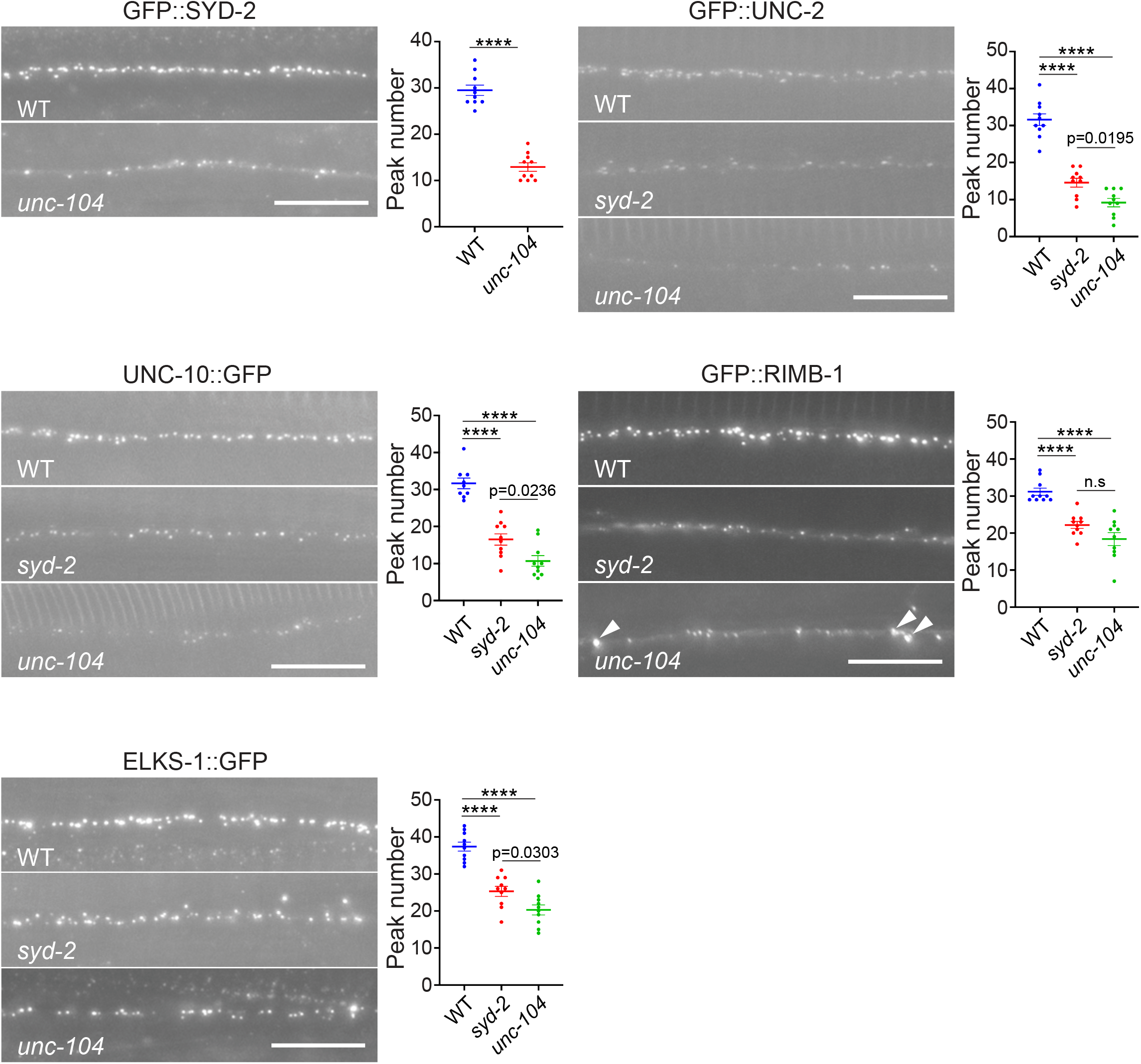
UNC-104/KIF1a is required for the presynaptic localization of UNC-2, UNC-10, RIMB-1, ELKS-1, and SYD-2. UNC-2, UNC-10, RIMB-1, and ELKS-1 in the dorsal nerve cords of wild type, *unc-104*, and *syd-2* mutant animals. SYD-2 in the dorsal nerve cords of wild type and *unc-104* mutant animals. Scale bar, 10 μm. The arrowheads indicate large aggregated RIMB-1 puncta. Error bars represent SEM. ****p<0.0001, for GFP::SYD-2, unpaired t-test and for the rest One-way ANOVA with Tukey’s post-hoc analysis.

## Discussion

In this study we used CRISPR/Cas9 genome editing to label endogenous CaV2/UNC-2 and performed a genetic screen to identify genes that control UNC-2 trafficking and localization. In addition to previously identified genes known to affect UNC-2 trafficking and function, we isolated *unc-10* and *syd-2*, two genes encoding key active zone proteins. Consistent with previous findings observed in knock-out mice of all RIM isoforms (Kaeser et al., 2011), *unc-10* null mutants exhibit a significant reduction of endogenous UNC-2 clusters at presynaptic active zones. Similarly, *syd-2* mutants also exhibit profound loss of presynaptic UNC-2 clusters in neurons. SYD-2 is known to have a broad function in the localization and stabilization of other active zone proteins at presynaptic terminals (Zhen and Jin, 1999; Dai et al., 2006; Patel et al., 2006). Here we show that in *syd-2* mutants, presynaptic UNC-10, RIMB-1, and ELKS-1 clusters were all reduced but not eliminated. Our genetic analysis indicates that the recruitment and stabilization of UNC-10 and RIMB-1 at presynaptic terminals by SYD-2 are critical for presynaptic UNC-2 clustering, while UNC-10 and ELKS-1 also contribute to presynaptic UNC-2 clustering in a SYD-2-independent manner.

Our data show that UNC-10 is the most consequential effector among active zone proteins in localizing UNC-2 to presynaptic terminals. A similar reduction in the levels of CaV2 channels was observed in RIM1 and RIM2 double KO mice, suggesting this is a conserved process (Kaeser et al., 2011). Previous studies determined that CaV2 channels bind to the PDZ domain of RIM proteins via their C terminal sequence (D/E-X-WC-COOH), which does not conform to classical PDZ binding motifs. While the *C. elegans* CaV2 ortholog, UNC-2, is divergent from this motif sequence, it matches the classical PDZ binding motif (X-*ϕ*-X-*ϕ*-COOH), suggesting a co-evolution of RIM and CaV2 proteins. A previous study using isothermal calorimetry showed that the RIM PDZ domain stoichiometrically interacts with the C-terminal sequences of CaV2 channels *in vitro* (Kaeser et al., 2011). Consistent with that study, our data showed that copy numbers of UNC-2 and UNC-10 at presynaptic terminals are comparable.

Our genetic screen and analysis in *C. elegans* showed that SYD-2 is important for UNC-2 synaptic clustering. SYD-2 was first identified as a critical mediator of synaptic vesicle cluster formation and synapse assembly in a *C. elegans* genetic screen (Zhen and Jin, 1999). It has since been established that Liprin-*α* /SYD-2 is required for normal presynaptic morphogenesis and synaptic vesicle docking in *Drosophila* and mouse neurons (Miller et al., 2005; Astigarraga et al., 2010; Spangler et al., 2013; Wong et al., 2018). The action of SYD-2 on presynaptic UNC-2 clustering is more complicated than that of UNC-10, as SYD-2 has broad effects on active zone assembly and maintenance. Interestingly, here we show that SYD-2 is not absolutely required for active zone formation, since key active zone proteins such as UNC-10, ELKS-1, and RIMB-1 are clustered at active zones in *syd-2* mutant animals, albeit at reduced levels. Indeed, previous ultrastructural analyses of neuromuscular synapses showed that *syd-2* mutant animals have fewer and smaller presynaptic dense projections associated with fewer docked SVs than those of wild-type animals, indicating that synapse number is reduced but not eliminated in the absence of SYD-2 (Kittelmann et al., 2013). A moderate reduction of UNC-10 puncta in *syd-2* mutants cannot fully account for reduced UNC-2 clusters in *syd-2* mutants, since the number of UNC-2 clusters is more severely reduced in *syd-2* mutants than in *unc-10* mutants. It is possible that SYD-2 acts on other active zone proteins that control UNC-2 clustering independently of UNC-10. Supporting this possibility, our data showed that SYD-2 partly acts on RIMB-1 to control UNC-2 clustering.

Unexpectedly, the *elks-1* mutation reduced UNC-2 clustering in the absence of SYD-2. In *C. elegans elks-1* mutants alone do not exhibit any obvious presynaptic phenotype (Deken et al., 2005). However, it was noted that ELKS-1 interacts with SYD-2 and is required for SYD-2 gain-of-function mutants to generate abnormally large dense projections (Kittelmann et al., 2013). A recent study has shown that SYD-2 and ELKS-1 undergo phase separation *in vivo* at the initial stage of synaptogenesis and their fluidity is critical for efficient incorporation of active zone proteins (McDonald et al., 2020). Thus, although *elks-1;syd-2* double mutants exhibit a dense projection structure comparable to *syd-2* single mutants (Kittelmann et al., 2013), it is possible that the absence of both SYD-2 and ELKS-1 may cause a defect in the synaptic assembly that prevents UNC-2 clustering at active zones. On the other hand, mammalian ELKS/CAST binds to *β*4 or *α* 1 subunit of CaV2 (Kiyonaka et al., 2012), and *Drosophila* Bruchpilot/ELKS associates with Cacophony/CaV2 (Kittel et al., 2006). This raises the possibility that, in the absence of SYD-2, ELKS-1 is one of the remaining links that directly support UNC-2 clustering via a physical interaction, and the absence of both SYD-2 and ELKS-1 severely compromises UNC-2 presynaptic clustering.

One intriguing question is why UNC-10 and SYD-2 were not identified in prior *C. elegans* genetic studies with an integrated transgene overexpressing GFP-tagged UNC-2 (Saheki and Bargmann, 2009; Kushibiki et al., 2019). Presynaptic CaV2 channels are known to interact with several active zone proteins via direct and indirect interactions (Hibino et al., 2002; Fouquet et al., 2009; Kaeser et al., 2011; Kiyonaka et al., 2012; Tong et al., 2017; Luo et al., 2020). We speculate that in the absence of UNC-10 or SYD-2, the residual overexpressed UNC-2 at presynaptic active zones due to the weaker redundant interactions with other active zone proteins shown here, may not allow a detectable reduction in signal, while endogenously tagged UNC-2 does. These types of redundant interactions are common amongst active zone proteins. It is well documented that while the loss of individual active zone components generally causes at best mild synaptic deficits, simultaneous loss of two components results in severe defects in active zone organization (Acuna et al., 2016; Wang et al., 2016; Kushibiki et al., 2019).

On the other hand, there is compelling evidence that an increase in certain active zone proteins upregulates CaV2 channel density and synaptic transmission efficiency. For example, in *Drosophila*, it has been reported that presynaptic homeostatic plasticity accompanies presynaptic changes in RBP/RIM-BP, Bruchpilot/ELKS, and Cacophony/CaV2 (Muller et al., 2012; Muller et al., 2015; Bohme et al., 2019; Gratz et al., 2019). In future studies, it will be interesting to investigate whether and how altered synaptic activity reorganizes the active zone and modifies UNC-2 channel density and synaptic transmission in *C. elegans*.

## Methods and Materials

### Genetic screen for mutants that have an altered UNC-2 localization

EMS mutagenized F2 progeny of *slo-1(eg142);unc-2(cim114[GFP::unc-2(L218S)])* animals were screened for 1 mM aldicarb resistance after 60 min. Based on their movement phenotypes, the confirmed candidate animals were tested for complementation with *unc-36(e250), unc-2(e55), calf-1(ky867)*, and *unc-10(md1117)*. As *unc-2(e55)* and *unc-10(md1117)* mutations were on X chromosome, we used extrachromosomally rescued transgenic animals to perform complementation testing. Select strains were subjected to whole-genome sequencing to identify specific mutation sites.

### CRISPR/Cas9 mediated genome editing

GFP or mScarlet was inserted into the genome using the co-conversion method (Arribere et al., 2014; Cheung et al., 2020). Genome editing was performed by injecting either plasmids expressing guide RNA and Cas9 or a riboprotein mixture of synthesized guide RNA (IDT Inc., IA, USA) and custom purified Cas9 protein into the gonad of young adult hermaphrodites. The 5’ and 3’ flanking sequences of genomic target insertion sites ranging from 500 bp to 1 kbp were subcloned to a plasmid containing the GFP or mScarlet coding sequence, and the resulting constructs were used as homologous repair templates.

### Aldicarb-induced paralysis assay

Age-matched adult animals (20 hr post L4) were assayed for their sensitivity to 1 mM aldicarb. Detailed protocol is published previously (Oh and Kim, 2017).

### Electrophysiology

Electrophysiological recordings from the *C. elegans* neuromuscular junction were performed as previously described (Richmond, 2006). Briefly, animals were immobilized with Histoacryl glue, and a lateral cuticle incision was made with a glass needle, exposing ventral neuromuscular junctions. Evoked postsynaptic currents were recorded from ventromedial body wall muscles, whole-cell voltage clamped at -60 mV in response to a 2ms depolarizing stimulus applied to the ventral nerve cord, using the following extracellular and intracellular solutions. The extracellular solution consisted of 150 mM NaCl, 5 mM KCl, 1 mM CaCl_2_, 4 mM MgCl_2_, 10 mM glucose, 5 mM sucrose, and 15 mM HEPES (pH 7.3, ∼340 mOsm. The patch pipette was filled with 120 mM KCl, 20 mM KOH, 4 mM MgCl_2_, 5 mM (N-tris[Hydroxymethyl] methyl-2-aminoethane-sulfonic acid), 0.25 mM CaCl_2_, 4 mM Na_2_ATP, 36 mM sucrose, and 5 mM EGTA (pH 7.2, ∼315 mOsm). Data were acquired using Pulse software (HEKA, Southboro, Massachusetts, United States) run on a Dell computer. Subsequent analysis and graphing were performed using Pulsefit (HEKA), Mini analysis (Synaptosoft Inc., Decatur, Georgia, United States) and Igor Pro (Wavemetrics, Lake Oswego, Oregon, United States).

### Microscopy and Image Analysis

Day one adult animals (20–24 hr post L4) were immobilized on a 2% agarose pad with a 6 mM levamisole solution in M9 buffer. Images were acquired using a 63x/1.4 numerical aperture on a Zeiss Axio-Observer Z1 microscope. Images were captured with an 84% quantum efficiency Zyla 4.2 plus (Oxford Instruments, RRID:SCR_017366) using a solid-state Spectra X light engine (Lumencor, OR, USA) as a light source in the same settings (light intensity and exposure time) for a given GFP transgenic animal. For consistent imaging of the same presynaptic terminals, we imaged an area of the dorsal nerve cord above the posterior gonad arm, which has minimal intestinal autofluorescence. This area consists of GABAergic and cholinergic presynaptic terminals and lacks cell bodies and dendrites. While we included wild-type controls to ensure that different results obtained in mutants were not due to changes in illumination, the results from different sessions are within the margin of error. The maximum projection was applied to images obtained as a Z-stack with a 0.2 μm interval. A line-scanning method in Metamorph (RRID:SCR_002368) was used for quantifying the average fluorescence intensity. In each image, a pixel intensity of 282 pixel length (30 μm) was measured, followed by a subtraction of background intensity from adjacent 282 pixels. From background-subtracted intensity values, peaks with a larger than a threshold value are identified as peaks of clusters. The threshold was arbitrarily set to exclude small background fluctuations and this same threshold value was applied to all the images of a given protein; one threshold value for all the GFP::UNC-2 images, one value for all the UNC-10, etc., except for Figure 7 where a different threshold was set because a higher light intensity was used to image GFP::UNC-2 in *syd-2(ok217)* mutant background animals.

### Microscopic relative copy number quantification of active zone proteins

To compare average and maximum puncta intensities between active zone proteins, we used a Prime 95B camera (Teledyne Photometrics, RRID:SCR_018464) to acquire Z-stack images from the dorsal nerve cord of transgenic animals with endogenously GFP-tagged active zone proteins or CaV2 UNC-2 channels. The high quantum efficiency (95%) and large pixel size (11 μm) of the camera allowed to image all of the tested active zone proteins and CaV2/UNC-2 in the same acquisition setting (light intensity and exposure time). To quantify puncta intensity, we first generated a maximum intensity projection from the Z-stack images, generated a binary mask derived from adaptive thresholding, applied the binary mask to a select dorsal cord region of the maximum intensity projection image, and then used Integrative Morphology Analysis (Metamorph 7.10) to get the average and maximum intensities of individual puncta. The final intensity values were obtained by subtracting a background intensity value of an adjacent area. The average and maximum intensities from 10 animals were pooled together for analysis.

### Statistical analysis

We used Prism 9 (RRID:SCR_002798) to perform statistical analysis. Sample size, p values, and statistical tests used presented in the figure legends. Sample size represents independent biological replicates. All of the raw data and statistical analysis summaries are included in the source data file.

---

**Figure 1-supplement 1.**
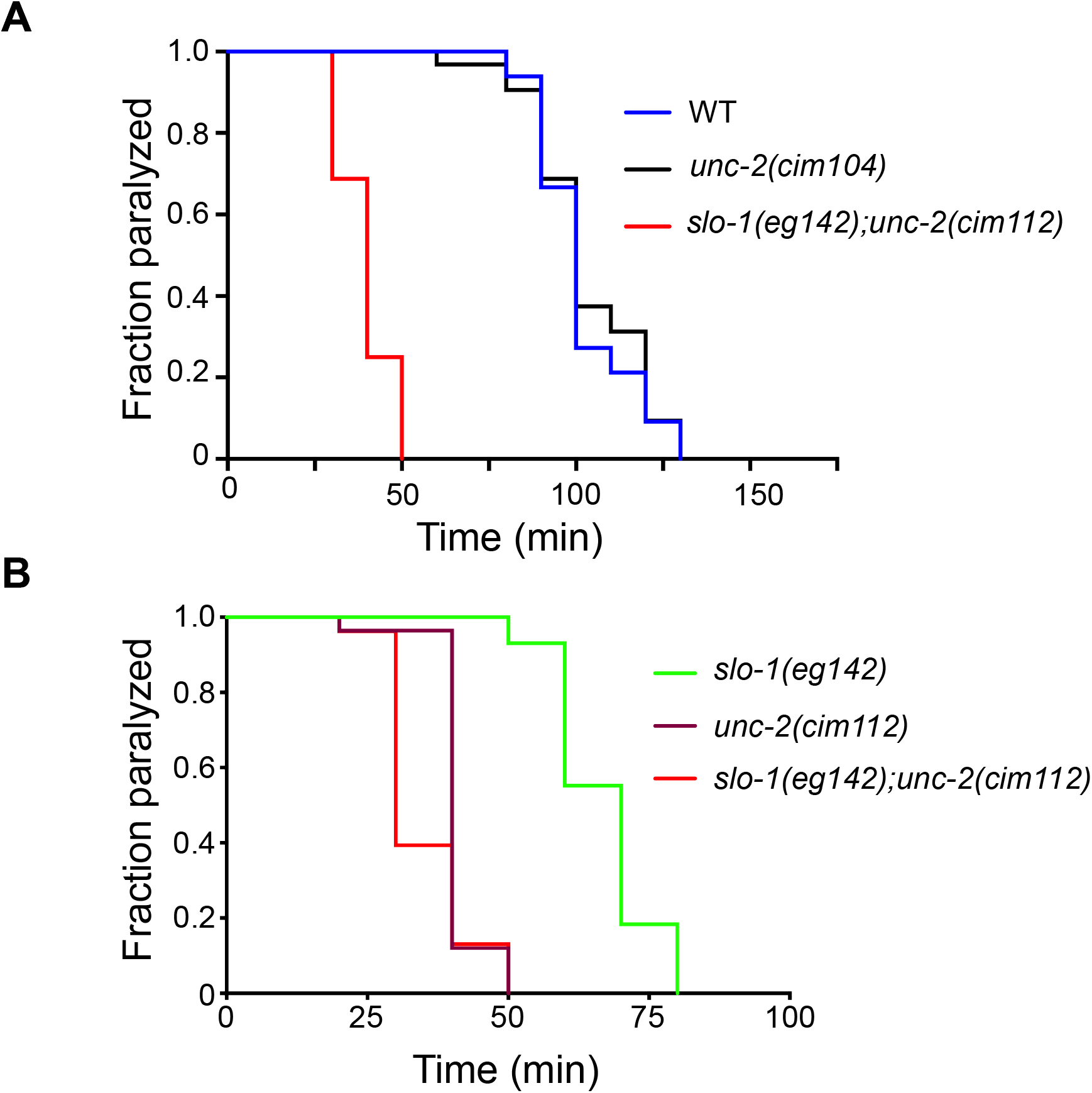
Aldicarb-mediated paralysis assays show that endogenous GFP tagging of UNC-2 does not alter synaptic function. (**A**) While the aldicarb sensitivity of *unc-2(cim104[GFP::unc-2])* animals is comparable to wild-type N2 animals, *slo-1(eg142);unc-2(cim112[GFP::unc-2S218L])* animals are hypersensitive. p < 0.0001, Log-rank survival test. (**B**) Comparison of *slo-1(eg142), unc-2(cim112[GFP::unc-2S218L])*, and *slo-1(eg142);unc-2(cim112)* animals in aldicarb sensitivity. p < 0.0001, Log-rank survival test.

**Figure 1-supplement 2.**
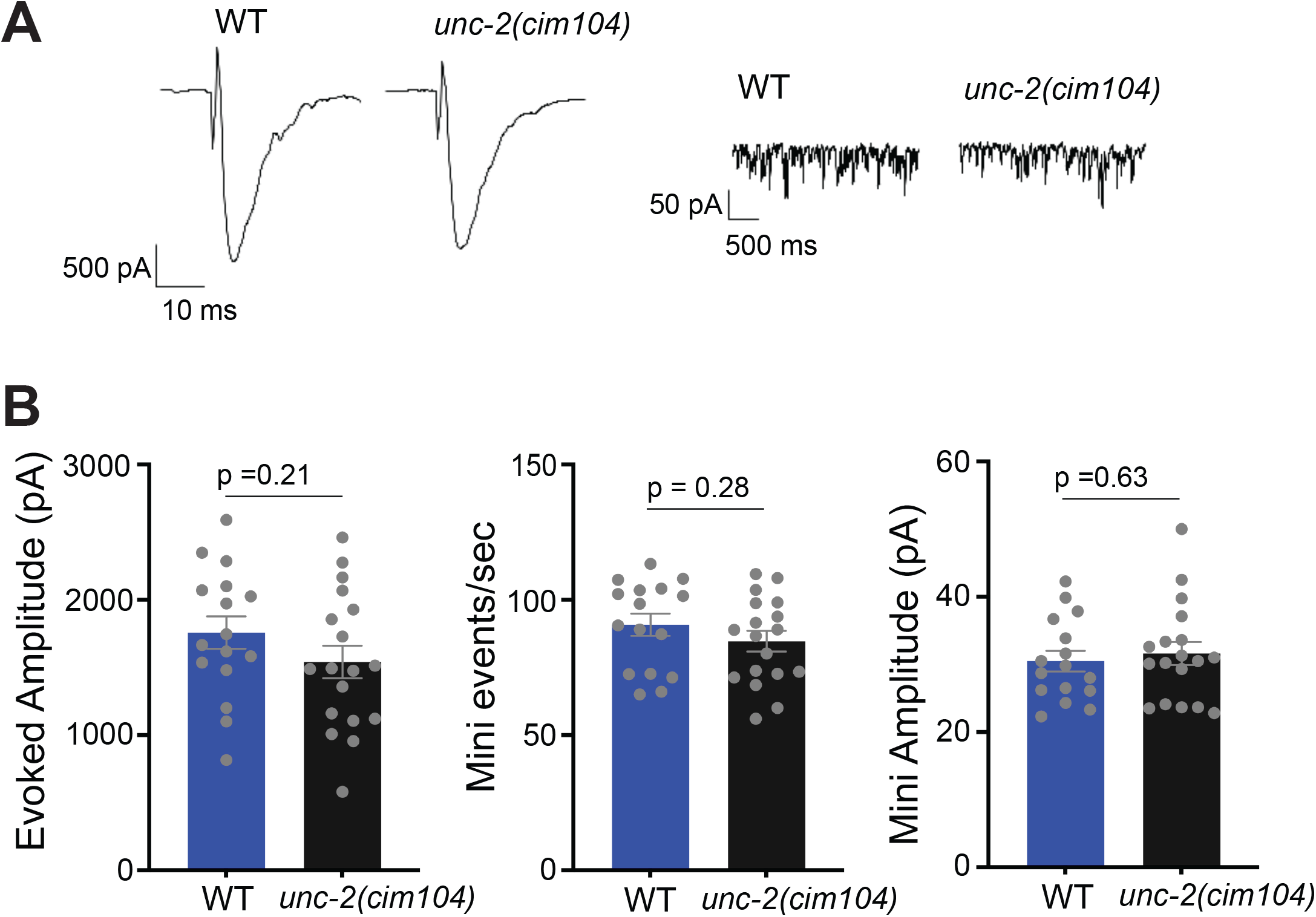
Electrophysiology at the neuromuscular junction show that endogenous GFP tagging of UNC-2 does not alter synaptic function. Patch clamp electrophysiology at muscle cells showed that *unc-2(cim104[GFP::unc-2])* animals exhibit a normal evoke response, mini frequency, and mini amplitude at 1 mM extracellular calcium ions in compared to wild-type animals. two-tailed t-test.

**Figure 1-supplement 3.**
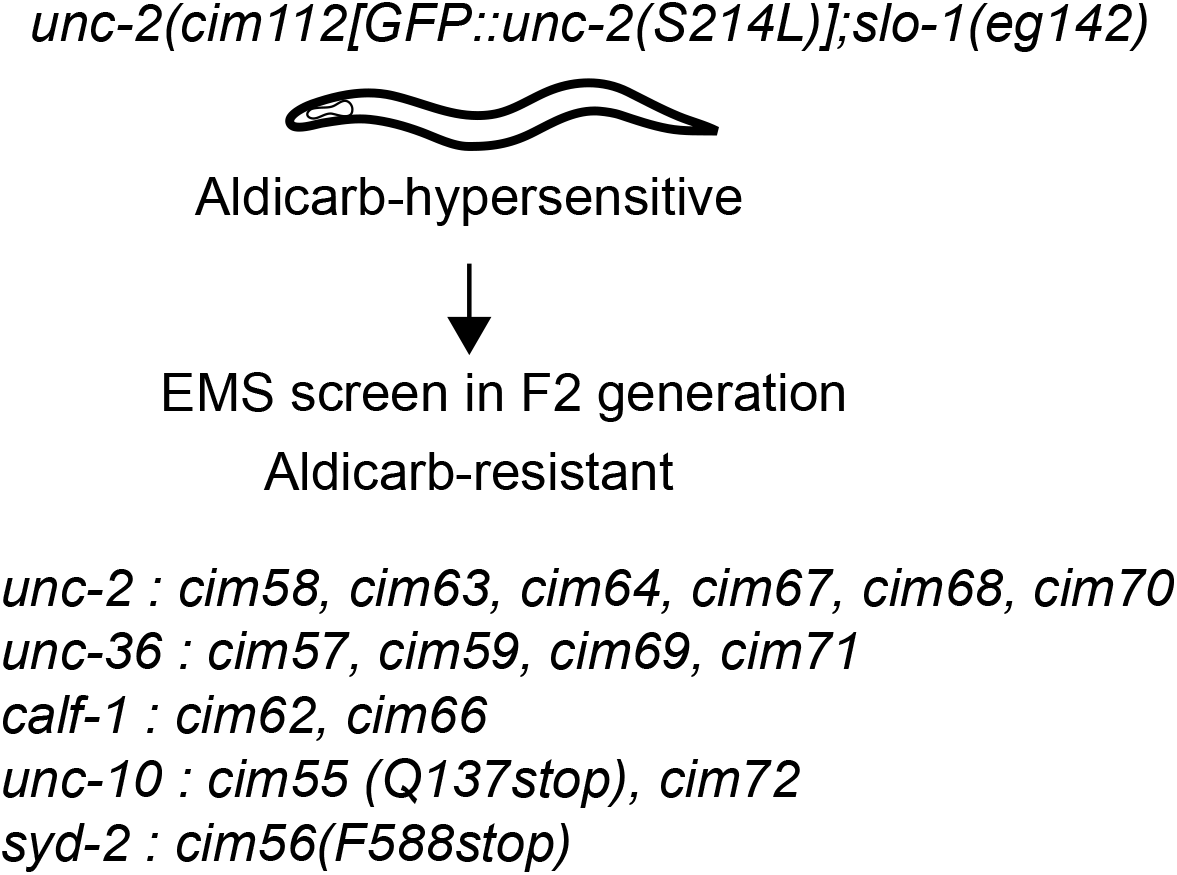
A genetic suppressor screen with *slo-1(eg142);unc-2(cim112[GFP::unc-2S218L])*. The screen with approximately 2500 haploid genome size yielded not only genes that affect UNC-2 function and trafficking, but also two genes that encode active zone proteins UNC-10/RIM and SYD-2/Liprin-*α*. Because we obtained two independent alleles of *calf-1*, which encodes a small protein with 177 amino acid residues, we consider that the screen is saturated or close to saturation.

**Figure 2-supplement 1.**
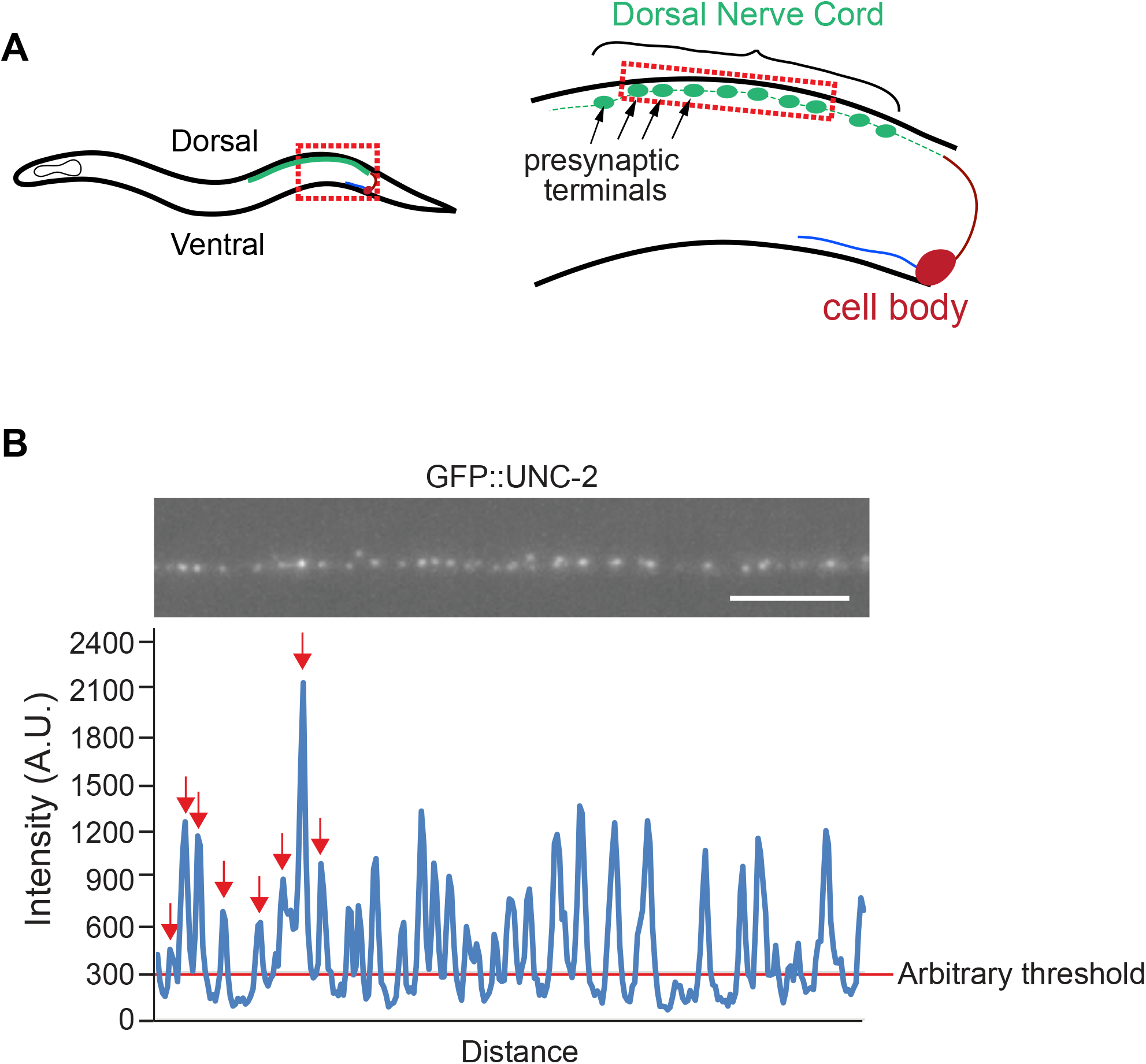
Schematics of the imaging strategy of the dorsal nerve cord. (**A**) The imaged area is part of the dorsal nerve cord positioned above the posterior gonad arm. This area consists of presynaptic terminals of GABAnergic and cholinergic neurons and lacks dendrites or cell bodies. (**B**) Peak intensity and peak number were calculated from the linescan method (see in Materials and Methods for details). Scale bar, 5 μm.

**Figure 7-supplement 1.**
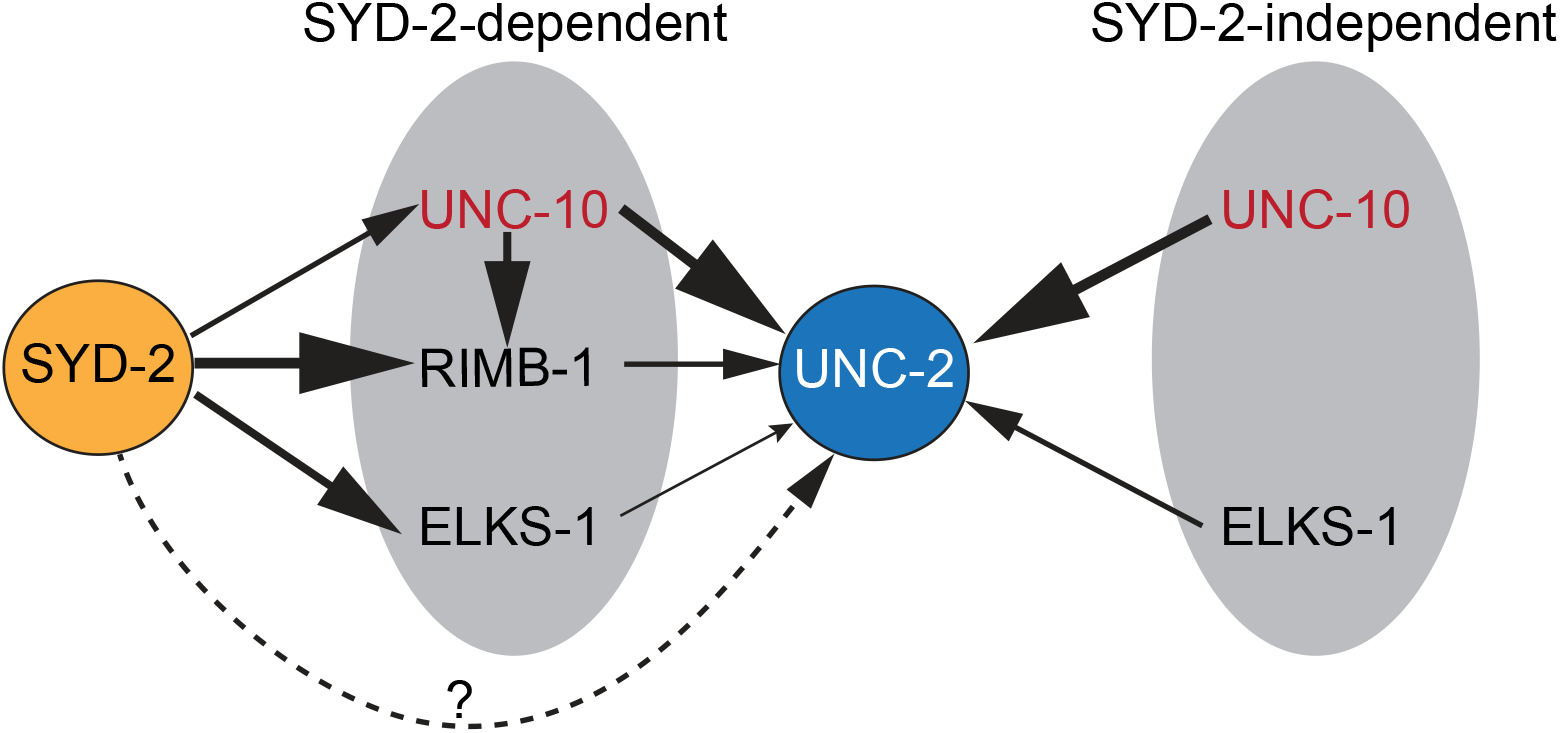
A genetic model for SYD-2-dependent and -independent mechanisms of presynaptic UNC-2 clustering. SYD-2 partly regulates presynaptic localization of UNC-10, RIMB-1, and ELKS-1. This regulation has some effects on UNC-2 clustering mainly through UNC-10 and RIMB-1. However, since a fraction of UNC-10 localizes to presynaptic terminals even in the absence of SYD-2, UNC-10, with a minor role of ELKS-1, mediates UNC-2 clustering independently of SYD-2.

**Video 1. Time lapse videos** of endogenously GFP-tagged RIMB-1 in the dorsal nerve cords of wild-type and *unc-104(e1265)* mutant animals.

